# A20 restriction of nitric oxide production restores macrophage bioenergetic balance

**DOI:** 10.1101/2025.10.26.684676

**Authors:** Mei-po Chan, Rommel Advincula, Philip Achacoso, Elizabeth A. Grossman, Daniel K. Nomura, Barbara A. Malynn, Averil Ma

## Abstract

Macrophages play critical roles in regulating host responses to microbial pathogens and other forms of tissue stress and can acquire pro-inflammatory or tissue reparative phenotypes. A20, or TNFAIP3, is a potent regulator of innate immune cell functions and is extensively linked to human inflammatory and autoimmune diseases. We now find that A20 is a powerful regulator of both glycolytic and mitochondrial respiration in macrophages. Differentiated macrophages that are acutely rendered A20 deficient exhibit increased glycolytic activity and markedly decreased mitochondrial respiration after LPS stimulation. These cells are unable to repolarize from an M1-like to M2-like phenotype. Compromised mitochondrial oxygen consumption in A20 deficient macrophages is caused by increased nitric oxide production. Inhibition or genetic ablation of inducible nitric oxide synthase (iNOS) restores mitochondrial oxidative phosphorylation and lactate production in these cells. These metabolic perturbations occur independently of exaggerated cytokine production and despite robust production of IL-10. Therefore, A20 prevents Warburg-like aerobic glycolysis and restores macrophage homeostasis.

## Introduction

Activation of immune cells is associated with a rapid burst in glycolytic activity along with mitochondrial remodeling (Buck et al., 2016). Several days post-activation, T cells return to more basal states associated with lower glycolysis and persistently increased mitochondrial respiration (Makowski et al., 2020). This latter transition may be important for maintaining an epigenetic state that retains physiological memory of past cellular exposures. It may also facilitate feedback mechanisms that control the duration and longitudinal character of immune responses.

Though less studied than T cells, macrophages appear to undergo similar metabolic changes during activation with ligands such as LPS (O’Neill and Pearce, 2016; Galván-Peña and O’Neill, 2014). Initial LPS stimulation of these cells triggers a glycolytic surge that provides energy for cytokine transcription, translation, and secretion. Subsequent subsiding of glycolysis with a relative shift back to mitochondrial oxidative phosphorylation coincides with transitions to relative LPS tolerance or to cells that express anti-inflammatory molecules associated with wound repair and tissue restoration, e.g., M2 macrophages. Importantly, while selected metabolic enzymes have been implicated in these processes (Mills et al., 2016), the cell biological mechanisms that couple immune cell activation with metabolic shifts are incompletely understood.

The ubiquitin editing protein A20, also known as TNF-α-induced protein 3 (TNFAIP3), is a susceptibility gene for multiple diseases including rheumatoid arthritis, Crohn’s disease, systemic lupus erythematosus, psoriasis, multiple sclerosis, scleroderma, and asthma (Ma and Malynn, 2012). In addition, patients with mono-allelic loss-of-function mutations of A20 develop mucosal and skin ulcers resembling Behçet’s disease as well as a growing number of other autoinflammatory and autoimmune disorders (Zhou et al., 2016; Ohnishi et al., 2017; Takagi et al., 2017; Aeschlimann et al., 2018). In parallel, A20 deficiency or mutations in mice leads to the development of spontaneous inflammation and autoimmune phenotypes (Boone et al., 2004; Lee et al., 2000; Tavares et al., 2010; Razani et al., 2020). Mice lacking A20 in selected innate immune cell types exhibit phenotypes that highlight the importance of A20 functions in macrophages, dendritic cells, and innate lymphoid cells (Hammer et al., 2011; Kool et al., 2011; Matmati et al., 2011; Schneider et al., 2018). A20 also regulates myelopoiesis, and perturbed myeloid differentiation may contribute to altered functions of mature cells (Muto et al., 2020). We have now explored the role of A20 in regulating metabolic transitions of macrophages, using a tamoxifen inducible system that allows myeloid cells to undergo normal hematopoiesis prior to A20 deletion. We find that A20 is a potent regulator of both glycolytic and mitochondrial activity in activated macrophages.

## Results and Discussion

### Induced deletion of A20 in macrophages promotes aerobic glycolysis

To understand how A20 regulates macrophage functions independently from developmental events, we interbred A20^Flox/Flox^ mice with mice bearing the ROSA/ER-Cre gene (Tavares et al., 2010). We generated bone marrow derived macrophages (BMDMs) from the resulting A20^Flox/Flox^ ROSA/ER-Cre+, A20^Flox/+^ ROSA/ER-Cre-, and A20^Flox/Flox^ ROSA/ER-Cre-mice, and established conditions for deleting A20 from A20^Flox/Flox^ ROSA/ER-Cre+ bone marrow cells in vitro. We call these cells A20^tiKO^ BMDMs. While Cre negative BMDMs induced expression of A20 protein within one hour after LPS stimulation, which gradually returned to basal levels over 16 hours, stimulation of A20^tiKO^ BMDMs expressed no detectable protein (**Fig. 1A**). LPS stimulation of A20^tiKO^ cells caused increased production of pro-inflammatory cytokines such as TNF and IL-6 when compared with control BMDMs, consistent with prior findings with A20^-/-^ BMDMs (**Fig. 1B**) (Boone et al., 2004; Turer et al., 2008). As macrophages develop tolerance to LPS and are thought to restore homeostatic functions after 16-48 hours (Seeley and Ghosh, 2017; Kotwal and Chien, 2017), we studied macrophages 24 hours after LPS stimulation. Remarkably, LPS stimulation of A20^tiKO^ BMDMs led to visibly changed media when compared to control cells (**Fig. 1C**). As culture medium acidification is most commonly caused by a change in lactate production, lactate levels in the culture supernatants were measured. A20^tiKO^ BMDMs produced markedly more lactate than control cells at 18 and 24 hours after LPS stimulation (**Fig. 1D**). This exaggerated lactate production was not observed after stimulation with other TLR ligands (**Fig. 1E**). FACS analyses revealed that cell survival was similar in A20^tiKO^ and WT BMDMs at 24 hours post LPS stimulation (**Fig. 1F**). As lactate is an end product of glycolysis, we measured glycolytic activity of these cells via Seahorse extracellular proton flux assay. Prior to LPS stimulation, both A20^WT^ and A20^tiKO^ BMDMs exhibited a similarly modest extracellular acidification rate (ECAR) (**Fig. 1G**). Consistent with prior studies (Galván-Peña and O’Neill, 2014), LPS caused increased glycolysis in normal macrophages (**Fig. 1G**). Importantly, after LPS stimulation, the ECAR of A20^tiKO^ cells was markedly higher than in LPS stimulated WT cells (**Fig. 1G**). After rotenone/antimycin A mediated inhibition of mitochondrial respiration, both A20^tiKO^ and WT cells enhanced proton efflux, with A20^tiKO^ cells showing greater glycolytic capacity (**Fig. 1G**). The differential ECAR between maximal glycolytic capacity and basal rate was slightly greater in WT cells, suggesting diminished glycolytic reserve in A20^tiKO^ cells (**Fig. 1G**).

**Figure 1.**
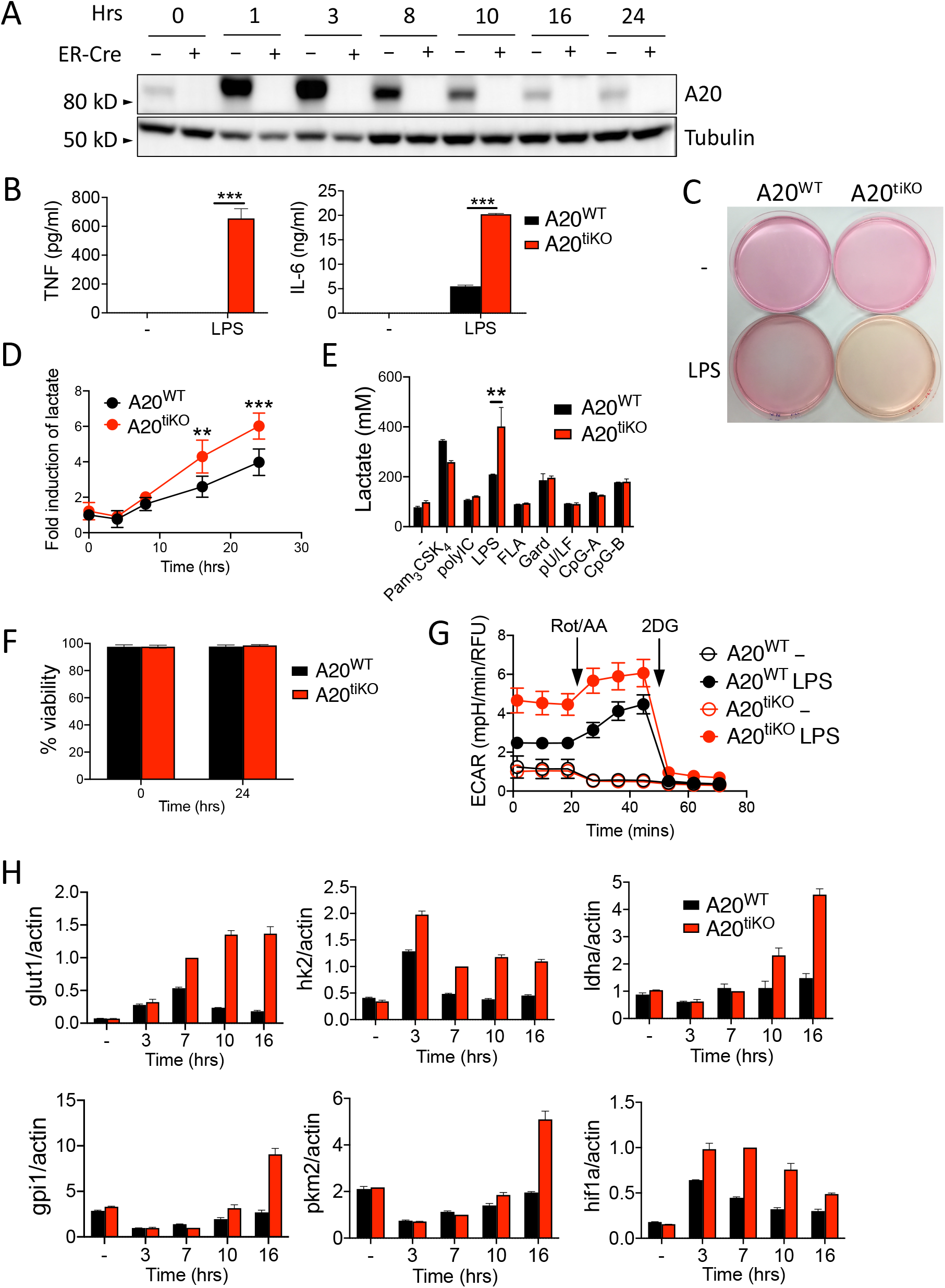
Inducible deletion of A20 from BMDMs leads to enhanced expression of glycolytic enzymes and glycolytic activity after LPS stimulation. **(A)** Immunoblot of A20 expression in A20^Flox/Flox^ Rosa/ER-Cre (“ER-Cre +” or A20^tiKO^) and A20^Flox/Flox^ (“ER-Cre –”) BMDMs after 4-OH-T mediated deletion of A20 followed by LPS stimulation for the indicated number of hours. Data are representative of three independent experiments. **(B)** ELISA of TNF and IL-6 secretion from A20^tiKO^ (red columns) and WT (black columns) BMDMs after LPS. Mean values ± SD are shown. *** indicates p<0.001 by Student’s T test. Data are representative of three independent experiments. **(C)** Gross images of culture media from A20^tiKO^ and WT BMDMs with or without LPS stimulation. Data are representative of at least three experiments. **(D)** Lactate secretion from BMDMs of indicated genotypes displayed as fold induction of LPS stimulated cells at indicated time points after LPS relative to unstimulated cells. Pooled results from six independent experiments are shown. Mean values ± SD are shown. ** indicates p<0.01 and *** indicates p<0.001 by one way ANOVA. **(E)** Lactate secretion from A20^tiKO^ (red columns) and WT (black columns) BMDMs after stimulation with indicated TLR ligands. Mean values ± SD are shown. ** indicates p<0.01 by Student’s T test. Data are representative of two independent experiments. **(F)** Flow cytometric quantification of BMDM cell viability via live-dead staining. Data are representative of two independent experiments. Mean values ± SD are shown. **(G)** Extracellular acidification rate (ECAR) of A20^tiKO^ and WT BMDMs with and without LPS stimulation analyzed by Seahorse proton flux assay. Rotenone/antimycin A and 2-deoxyglucose (2-DG) added at indicated times. Mean values ± SD are shown. Data are representative of three independent experiments. **(H)** Real-time quantitative PCR analyses of *glut1, hk2, ldha, gpi1, pkm2* and *hif1a* mRNA expression relative to *actin* mRNA in indicated genotypes of cells at indicated times after LPS stimulation. Mean values ± SD are shown. Data are representative of two independent experiments. All TLR stimulations were conducted for 24 hours unless otherwise stated.

The glycolytic surge in activated lymphocytes is supported by multiple molecular adaptations. Glut1 is a glucose transporter which is transcriptionally induced in CD4^+^ T cells, and is essential for their activation and differentiation into effector cells (Macintyre et al., 2014). Hexokinase2 phosphorylates intracellular glucose, preventing its transport out of the cell, and represents the first step of glycolysis. Lactate dehydrogenase (LDH) converts pyruvate to lactate and regenerates nicotinamide adenine dinucleotide (NAD) to support glycolysis. Glucose 6 phosphate isomerase (GPI) converts glucose-6-phosphate to fructose-6-phosphate and represents the second step of glycolysis. Pyruvate kinase M2 (PKM2) converts phosphoenolpyruvate to pyruvate. HIF1α is a transcription factor that drives expression of several glycolytic enzymes. After LPS stimulation, expression of mRNAs for all these molecules was markedly increased in A20 deficient macrophages when compared to control cells (**Fig. 1H**). Hence, A20 restrains multiple facets of glycolysis that collaborate to limit the glycolytic surge accompanying LPS stimulation. This increased glycolytic activity may support enhanced production and secretion of cytokines by A20 deficient macrophages.

### Induced deletion of A20 in macrophages suppresses mitochondrial respiration and prevents M2 repolarization after LPS

After LPS stimulation, macrophages can assume tissue reparative functions, reduction of glycolysis, and a return to predominantly mitochondrial oxidative phosphorylation for ATP production. We thus assayed mitochondrial respiration by measuring the oxygen consumption rate (OCR) of A20^tiKO^ and control BMDMs via Seahorse assays. Consistent with prior work, the OCR of WT BMDMs increased after LPS stimulation (**Fig. 2A**). By contrast, A20^tiKO^ BMDMs, which contained mildly elevated levels of OCR when compared to unstimulated WT cells, exhibited markedly decreased OCR after LPS stimulation (**Fig. 2A**, see basal OCR). Oligomycin inhibits ATP synthase and the OCR in oligomycin treated cells reflects the amount of ATP generated by the mitochondrial electron transport chain. A20^tiKO^ BMDMs exhibited negligible decrements of OCR after oligomycin (**Fig. 2A**, see ATP production). FCCP uncouples the proton gradient generated by the electron transport chain, unveiling the maximal oxygen consumption potential of mitochondria in normal cells. While WT cells and unstimulated A20^tiKO^ BMDMs increased their OCR to maximal levels, LPS stimulated A20^tiKO^ BMDMs completely failed to increase their OCR in response to FCCP (**Fig. 2A**, see maximal). Therefore, LPS stimulation cripples the ability of mitochondrial electron transport complexes in A20^tiKO^ BMDMs to consume oxygen even when these complexes are uncoupled from their electron transport functions. The difference between FCCP induced maximal OCR and basal OCR, termed the spare respiratory capacity (SRC), reflects the cell’s ability to marshal reserves for energy production and is an indicator of cellular fitness (**Fig. 2A**, see SRC). In this regard, A20^tiKO^ BMDMs displayed negligible SRC. Finally, inhibition of complexes I and III with rotenone and antimycin A blocks mitochondrial respiration and unveils oxygen consumption from non-mitochondrial sources. These treatments revealed modestly increased OCR in A20^tiKO^ BMDMs, consistent with the increased glycolysis observed in these cells. These respiratory defects in A20^tiKO^ BMDMs were triggered by LPS stimulation but not by other TLR ligands such as Pam_3_CSK_4_ and polyI:C (**Figs. 2B, 2C**). Hence, while A20^tiKO^ BMDMs conduct increased glycolysis after LPS stimulation, these cells perform markedly less mitochondrial respiration, thereby exhibiting a profound bioenergetic bias toward aerobic glycolysis. This dramatic metabolic shift from oxidative phosphorylation to aerobic glycolysis in A20 deficient macrophages constitutes a potent version of the Warburg effect initially described in cancer cells. Importantly, as A20 is a tumor suppressor (Schmitz et al., 2009; Kato et al., 2009; Compagno et al., 2009), the loss of this protein could drive aerobic glycolysis in some cancer cells as well.

**Figure 2.**
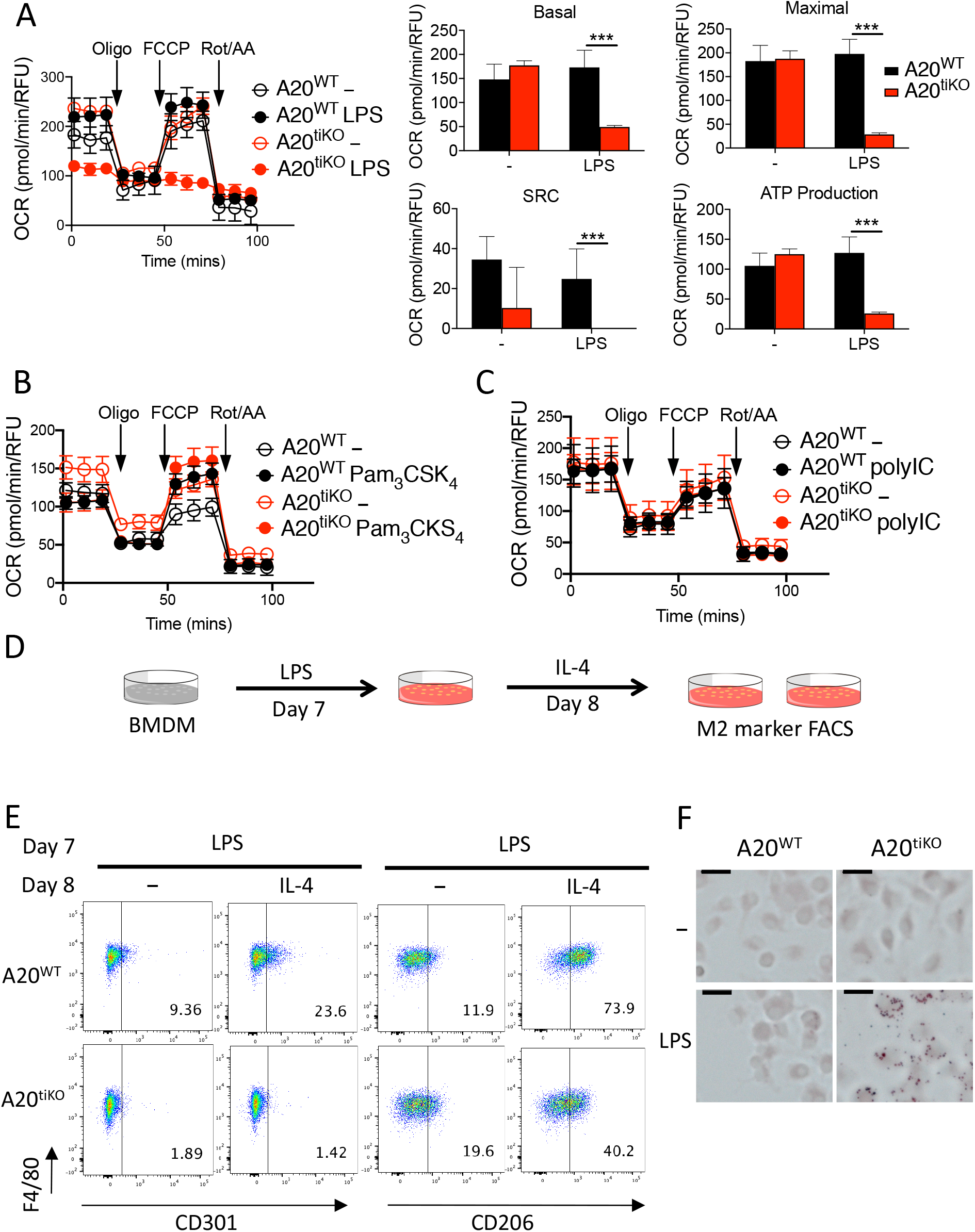
LPS activation cripples mitochondrial respiration and prevents M2 repolarization in A20^tiK O^ BMDMs. **(A)** Oxygen consumption rate (OCR) measured by Seahorse flux assay of LPS stimulated A20 deficient (red circles) and WT (black circles) BMDMs. Open circles represent unstimulated cells; Closed circles represent LPS stimulated cells. Mitochondrial stress tests with oligomycin (Oligo), FCCP, and rotenone/antimycin A injected at indicated times (arrows). Calculated levels of basal and maximal mitochondrial respiration, spare respiratory capacity (SRC) and ATP production shown at right. Mean values ± SD are shown. *** indicates p<0.001 by unpaired two-tailed T test. Data are representative of at least five independent experiments. **(B and C)** Mitochondrial stress tests of A20WT and A20^tiKO^ BMDMs after stimulation with Pam_3_CSK_4_ **(B)** or polyI:C **(C)**. Mean values ± SD are shown. Data are representative of three independent experiments. **(D)** Schematic representation of experimental design of M2 repolarization assay. On day 7 of BMDM differentiation, cells were treated with LPS for 24 hours, washed and stimulated with IL-4 or medium for another 24 hours. Flow cytometric measurement of M2 surface markers were assayed on day 9. **(E)** Representative flow cytometric analyses of CD301 and CD206 cell surface expression (M2 markers) on A20^tiKO^ and WT BMDMs after IL-4 mediated repolarization as described in (D). Mean values ± SD are shown. Data are representative of two independent experiments. **(F)** Oil red O staining of indicated genotypes of BMDMs after stimulation with LPS. Note lipid droplets inside A20^tiKO^ BMDMs. Scale bars represent 2 µm. Data are representative of three independent experiments.

The major source of ATP in M2 macrophages derives from oxidative phosphorylation by mitochondria (Jha et al., 2015; Vats et al., 2006). Hence, A20’s preservation of macrophage mitochondrial function might support M2 repolarization after LPS stimulation. We tested the ability of LPS stimulated A20^tiKO^ BMDMs to repolarize to an M2 phenotype after IL-4 stimulation. We stimulated A20^tiKO^ and A20^WT^ BMDMs with LPS for 24 hours, treated these cells with IL-4 for another 24 hours, and finally analyzed the expression of M2 markers by FACS (**Fig. 2D**). A20^tiKO^ BMDMs were largely unable to express the M2 markers CD301 and CD206 after repolarization with IL-4 (**Fig. 2E**). Instead, A20^tiKO^ BMDMs were strongly stained with Oil Red O, indicating enhanced lipid droplet formation and accumulation after LPS stimulation (**Fig. 2F**), a hallmark of inflammatory M1 macrophages (Rosas-Ballina et al., 2020; Wen et al., 2011). Such lipid droplets have been described as sites of eicosanoid synthesis, enzyme localization and cytokine storage in inflammatory leukocytes (Bozza and Viola, 2010; Jarc and Petan, 2020). Hence, A20 is required for the ability of activated macrophages to express type 2 markers and potentially evolve into cells supporting tissue repair. The persistence of M1 associated proteins likely contributes to the ability of these cells to drive inflammatory diseases.

### Mitochondria in A20 deficient macrophages exhibit defective membrane potential and compromised turnover

Healthy mitochondria generate an electrochemical proton gradient that fuels the electron transport chain and ATP generation. To better understand how A20 maintains mitochondrial respiration in LPS stimulated macrophages, we first measured the mitochondrial membrane potential in A20^WT^ and A20^tiKO^ cells. Consistent with our Seahorse assays (**Fig. 2A**), A20^tiKO^ BMDMs exhibited decreased transmembrane potential after LPS stimulation (**Fig. 3A**). This decreased membrane potential was not associated with reduced mitochondrial mass, as the ratio of mitochondrial DNA to nuclear DNA was preserved in A20^tiKO^ BMDMs (**Fig. 3B**). Immunoblotting analyses of the mitochondrial protein Tomm20 revealed normal amounts of this protein in both unstimulated and LPS stimulated A20^tiKO^ BMDMs, reinforcing the notion that mitochondrial mass is preserved in these cells (**Fig. 3C**). Taken together, these results indicate that mitochondria are structurally intact, but functionally compromised in LPS stimulated A20^tiKO^ BMDMs.

**Figure 3.**
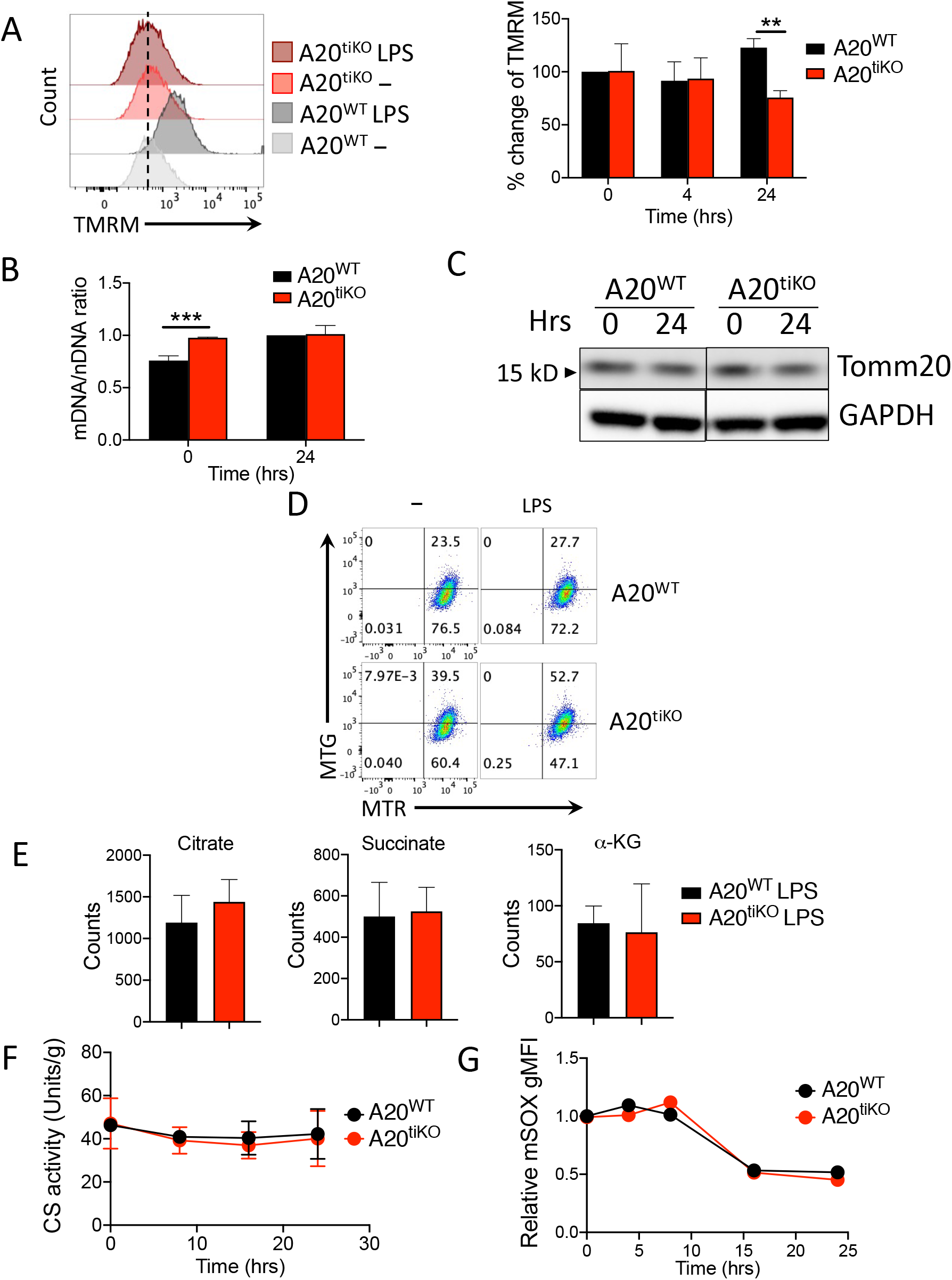
Mitochondria in A20^tiKO^ BMDMs exhibit lower membrane potential and decreased turnover. **(A)** FACS mediated measurement of mitochondrial transmembrane potential using tetramethylrhodamine methyl ester (TMRM) labeling. Representative histograms of fluorescent TMRM expression in BMDMs of the indicated genotypes shown at left. Percentage change in TMRM staining calculated at right from three independent experiments. Mean values ± SD are shown. ** indicates p< 0.01 by unpaired T test. **(B)** Real time qPCR analyses of mitochondrial and nuclear DNA from the indicated BMDMs. Mean values ± SD are shown. *** indicates p<0.001. Data are representative of two independent experiments. **(C)** Immunoblot analysis of mitochondrial Tomm20 protein expression in indicated BMDMs. GAPDH expression shown below as loading control. Data are representative of three independent experiments. **(D)** FACS analyses of mitochondrial turnover using differential labeling. BMDMs were first labeled with MitoTracker Green FM (MTG), then stimulated with LPS, and subsequently labeled with MitoTracker Red (MTR) prior to FACS analysis. In this format, MTG represent pre-existing (old) mitochondria; MTR represent all mitochondria present after LPS stimulation. Note that A20tiKO cells retain a higher proportion of MTG positive (old) mitochondria than WT cells, particularly after LPS stimulation (52.7% vs. 27.7%). Data are representative of three independent experiments. **(E)** Mass spectrometric quantification of indicated tricarboxylic acid (TCA) cycle intermediates in A20^tiKO^ (red) and WT (black) BMDMs after LPS stimulation. Counts reflect average of five replicates. **(F)** Fluorometric assay of citrate synthase (CS) activity of indicated cells after LPS stimulation was measured. **(G)** FACS analyses of mitochondrial superoxide production in A20^tiKO^ and WT BMDMs using using mitoSOX dye (mSOX). Data are representative of three independent experiments.

During metabolically demanding cellular activation events, damaged mitochondria undergo conformational changes and turnover to preserve the quantity and quality of these organelles (Gkikas et al., 2018). We thus assessed mitochondrial turnover in A20^WT^ and A20^tiKO^ BMDMs. We utilized differential labeling of mitochondria using MitoTracker Green (MTG) and MitoTracker Red (MTR) before and after LPS stimulation, to quantitate pre-existing and newly generated mitochondria. A20^tiKO^ BMDMs contained a larger fraction of older, pre-existing (MTG+) mitochondria than WT BMDMs without LPS, suggesting that basal mitochondrial turnover is compromised in A20^tiKO^ cells (**Fig. 3D**). This difference became much more pronounced after LPS stimulation, indicating that A20 may be particularly important for replacing senescent mitochondrial in activated macrophages (**Fig. 3D**). A20 dependent mitochondrial replacement thus appears to collaborate with A20’s ability to preserve mitochondrial OCR in a coordinated effort to preserve cellular energetic homeostasis.

To further investigate the defective mitochondrial respiration in A20 deficient macrophages, we performed targeted metabolomic studies on LPS stimulated cells. These assays revealed that A20^tiKO^ BMDMs contain similar levels of the tricarboxylic acid (TCA) cycle intermediates, succinate, citrate and α-ketoglutarate (α-KG) as control cells, suggesting that TCA cycling proceeds at sufficient rates in these cells to generate electron donors and substrates to the mitochondrial electron transport chain (**Fig. 3E**). We also directly assayed the activity of citrate synthase, the enzyme catalyzing the condensation between oxaloacetate and acetyl Co-A to generate citrate. This enzyme is considered the first step of the TCA cycle. These tests showed that citrate synthase activity was preserved in A20^tiKO^ BMDMs throughout 24 hours of LPS stimulation (**Fig 3F**). As the levels of citrate and succinate were comparable in WT and A20 deficient macrophages, previously described inhibition of these TCA cycle steps do not appear to depend upon A20 (Tannahill et al., 2013; Liu et al., 2017; Jha et al., 2015). Finally, as increased release of mitochondrial superoxide can cause dysfunctional mitochondria with augmented electron leaks, we measured oxidation of the superoxide indicator, MitoSOX (**Fig. 3G**). During the 24 hours of LPS stimulation, MitoSOX levels remained similar between A20^WT^ and A20^tiKO^ cells, implying that impaired mitochondrial respiration in A20^tiKO^ cells was not caused by excessive induction of mitochondrial superoxide. Taken together, these data suggest that acute A20 deficiency in macrophages causes defective mitochondrial respiration despite the availability of comparable TCA substrates and structurally intact mitochondria.

### A20 deficient macrophages accumulate increased reactive oxygen and nitrogen species, and exaggerated inducible nitric oxide synthase (iNOS) expression compromises mitochondrial respiration

Elevated cellular reactive oxygen species (ROS) and reactive nitrogen species (RNS) can compromise mitochondrial function by inhibiting electron transport chain enzymes (Kaludercic and Giorgio, 2016; Kowalska et al., 2020). Our metabolomic analyses revealed markedly decreased glutathione (GSH) levels in LPS stimulated A20^tiKO^ BMDMs (**Fig. 4A**). In addition, glycine, an essential and rate-limiting precursor of GSH, was also significantly lower in these BMDMs (**Fig. 4A**). As GSH is a major cellular antioxidant that reacts with and scavenges potentially deleterious ROS and RNS, we hypothesized that A20 deficient macrophages might accumulate excessive amounts of these reactive species. We measured cellular ROS/RNS levels using the fluorescent indicator dichlorodihydrofluorescein diacetate (DCFDA), and found that A20^tiKO^ BMDMs exhibited significantly higher levels of DCFDA staining after LPS stimulation (**Fig. 4B**). Among RNS, nitric oxide (NO) and its metabolite peroxynitrite inhibit electron transport chain enzymes by binding to O_2_ binding sites on these proteins (Stewart and Heales, 2003). We assayed NO levels and found that A20^tiKO^ BMDMs produced markedly more NO than control cells after LPS (**Fig. 4C**). This exaggerated NO production was not observed after stimulation with other TLR ligands (**Fig. 4C**). We next tested whether abrogating production of these species might restore mitochondrial defects. While allopurinol, a xanthine oxidase inhibitor, did not rescue OCR defects in LPS stimulated A20^tiKO^ BMDMs (**Fig. 4D**), the iNOS inhibitor 1400w effectively restored mitochondrial respiration (**Fig. 4E**) and M2 repolarization (Suppl. Fig. 1).

**Figure 4.**
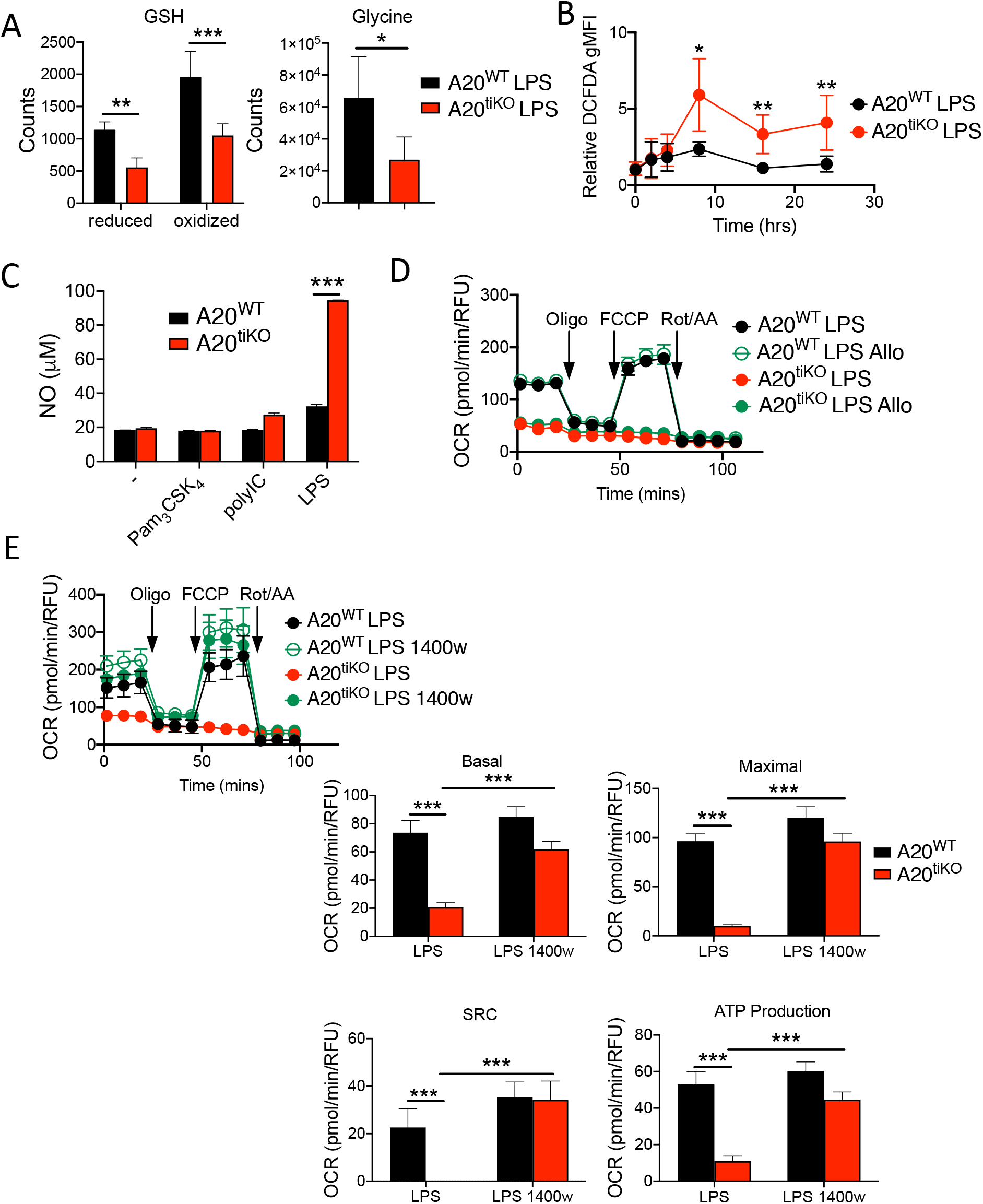
A20^tiKO^ BMDMs produce exaggerated amounts of nitric oxide (NO), and impaired mitochondrial respiration is rescued by inhibition of nitric oxide synthase (iNOS). **(A)** Mass spectrometric quantification of reduced and oxidized forms of glutathione (GSH) (left graph) and glycine (right graph) in A20^tiKO^ (red) and WT (black) BMDMs after LPS stimulation. * indicates p<0.05, ** indicates p<0.01, *** indicates p<0.001. Data are representative of 5 independent samples. **(B)** Flow cytometric analyses of cytosolic ROS/RNS species in LPS treated BMDMs of the indicated genotypes using dichlorodihydrofluorescein diacetate (DCFDA) dye. Y-axis represents geometric mean fluorescence intensity (gMFI) of each sample relative to untreated cells at the indicated timepoints after LPS stimulation. * indicates p<0.05, ** indicates p<0.01. Mean values ± SD are shown. Pooled results from five independent experiments are shown. **(C)** Calorimetric determination of nitric oxide (NO) production from A20^tiKO^ (red) and WT (black) BMDMs after stimulation with the indicated TLR ligands. Mean values ± SD are shown. *** indicates p<0.001. Data are representative of three independent experiments. **(D)** Seahorse mitochondrial stress test of A20^tiKO^ (red) and WT (black) BMDMs with or without xanthine oxidase inhibitor allopurinol (allo). Mean values ± SD are shown. Data are representative of three independent experiments. **(E)** Seahorse mitochondrial stress test of A20^tiKO^ (red) and WT (black) BMDMs with or without iNOS inhibitor 1400w. Basal and maximal respiratory rates, spare respiratory capacity (SRC), and ATP production are shown at right. Mean values ± SD are shown. *** indicates p<0.001. Data are representative of three independent experiments. In all panels, * indicates p<0.05, ** indicates p<0.01, *** indicates p<0.001 using unpaired two-tailed t test.

NO is produced by several nitric oxide synthase enzymes that are expressed in multiple cell types. To understand why A20^tiKO^ BMDMs secreted excessive amount of NO after LPS stimulation, we measured the expression of iNOS (*nos2*), the inducible isoform of NOS that is robustly expressed in macrophages and generates NO in a Ca^++^ independent fashion (Förstermann and Sessa, 2012). While resting BMDMs express negligible levels of iNOS mRNA and protein, these species were both expressed at markedly higher levels in A20^tiKO^ BMDMs than control cells after LPS stimulation (**Figs. 5A, B**). To define the contribution of iNOS to mitochondrial dysfunction in A20^tiKO^ BMDMs, we interbred A20^Flox/Flox^ ROSA/ER-Cre mice with iNOS^-/-^ mice (Laubach et al., 1995) and tested LPS responses of the compound mutant macrophages derived from these mice. Immunoblot analyses of A20 and iNOS expression confirmed elimination of these proteins in the appropriate genotypes (**Suppl. Fig. 2**). Stimulation of A20^tiKO^ iNOS^-/-^ BMDMs with LPS resulted in normalization of cellular ROS/RNS levels when compared with A20^tiKO^ BMDMs (**Fig. 5C**). Hence, iNOS is predominantly responsible for the exaggerated ROS/RNS species generated in LPS stimulated A20^tiKO^ BMDMs. Similar results were obtained with the iNOS inhibitor 1400w (**Suppl. Fig. 3**). Remarkably, extracellular flux assays revealed that ablation of iNOS completely restored mitochondrial respiration in A20^tiKO^ BMDMs (**Fig. 5D**). These A20^tiKO^ iNOS^-/-^ BMDMs also exhibited normal levels of ATP production and spare respiratory capacity (**Fig. 5D**). In addition to restoring deficient mitochondrial respiration, abrogation of iNOS expression also normalized the exaggerated lactate production observed in A20^tiKO^ BMDMs (**Fig. 5E**). Therefore, iNOS expression is a dominant mechanism regulating the balance between glycolytic and mitochondrial respiration.

**Figure 5.**
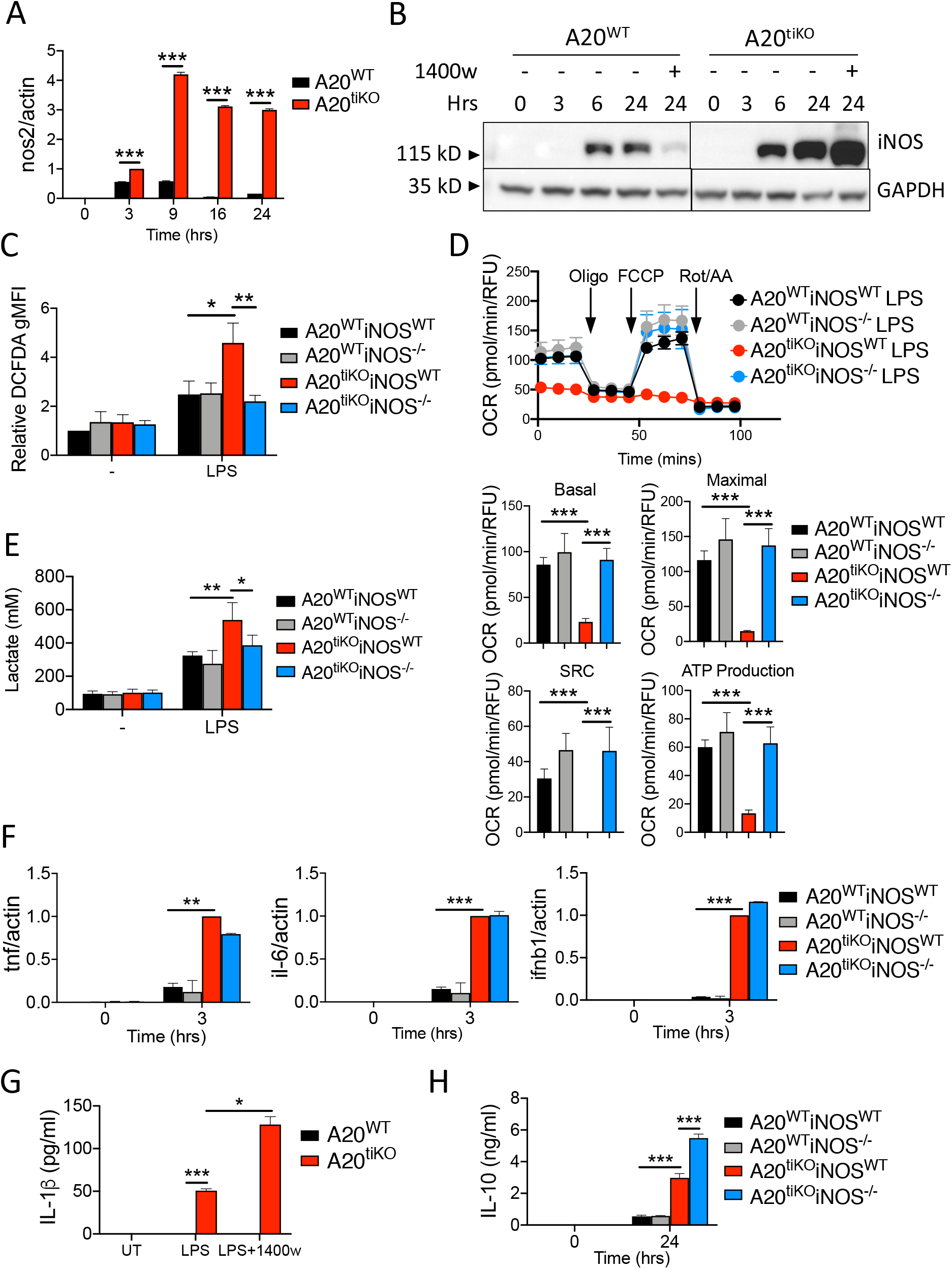
Expression of iNOS is greatly induced in A20^tiKO^ BMDMs, and genetic ablation of nos2 restores mitochondrial bioenergetic balance. **(A)** Real-time qPCR analyses of nos2 mRNA expression relative to actin mRNA in A20^tiKO^ (red) and A20^WT^ (black) BMDMs at indicated timepoints after LPS stimulation. Mean values ± SD are shown. *** indicates p<0.001 by unpaired T test. Data are representative of three independent experiments. **(B)** Immunoblot analyses of iNOS protein expressions in indicated BMDMs at indicated timepoints after LPS stimulation. GAPDH protein expression shown below as loading control. The iNOS inhibitor 1400w inhibits iNOS activity rather than iNOS expression and is shown as a control. Data are representative of three independent experiments. **(C)** Flow cytometry analysis of ROS/RNS species by DCFDA staining in the indicated genotypes of BMDMs after LPS stimulation. DCFDA staining intensity is plotted relative to unstimulated cells. Note that increased ROS/RNS in LPS stimulated A20^tiKO^ cells (red) is normalized in LPS stimulated A20^tiKO^ iNOS^-/-^ cells (blue). Mean values ± SD are shown. * indicates p<0.05, ** indicates p<0.01. Data are representative of three independent experiments. **(D)** Seahorse mitochondrial stress test of BMDMs from the indicated genotypes of mice. Note that depressed basal and maximal oxygen consumption rates (OCR) in A20^tiKO^ cells (red) is normalized in LPS stimulated A20^tiKO^ iNOS^-/-^ cells (blue). Graphic representation of basal and maximal OCRs, spare respiratory capacity (SRC), and ATP production are shown below. Mean values ± SD are shown. *** indicates p<0.001. Data are representative of four independent experiments. **(E)** Colorimetric measurements of lactate secretion by indicated genotypes of cells. Mean values ± SD are shown. * indicates p< 0.05, ** indicates p<0.01 by unpaired T test. Data are representative of three independent experiments. **(F)** Real-time quantitative PCR analyses of *tnf, il-6 and ifnb1* mRNA expression relative to *actin* in the respective BMDMs after LPS stimulation. Mean values ± SD are shown. Data are representative of three independent experiments. **(G)** ELISA of IL-1β secretion by A20^WT^ and A20^tiKO^ and WT BMDMs after indicated treatments. Mean values ± SD are shown. * indicates p<0.05, *** indicates p<0.001. Data are representative of two independent experiments. **(H)** ELISA of IL-10 secretion by A20^WT^ and A20^tiKO^ and WT BMDMs after indicated treatments. Mean values ± SD are shown. *** indicates p<0.001 by unpaired T test. Data are representative of two independent experiments.

### Resuscitation of mitochondrial respiration in A20^tiKO^ iNOS^-/-^ BMDMs occurs independently from A20 regulated inflammatory cytokines and inflammasome activation

A20 deficient macrophage express greater amounts of inflammatory cytokines than control cells in response to LPS stimulation (Boone et al., 2004; Turer et al., 2008). These cytokines could theoretically contribute to mitochondrial dysfunction in A20^tiKO^ BMDMs. We tested cytokine responses in A20^tiKO^ and A20^tiKO^ iNOS^-/-^ BMDMs after LPS stimulation and found that exaggerated production of TNF, IL-6, and IFN-β mRNAs by A20^tiKO^ BMDMs was not abrogated by elimination of iNOS expression in A20^tiKO^ iNOS^-/-^ BMDMs (**Fig. 5F**). Therefore, iNOS deficiency decouples mitochondrial dysfunction in A20^tiKO^ BMDMs from exaggerated secretion of pro-inflammatory cytokines.

ROS have been described to mediate a second step to activate NLRP3 inflammasomes (Zhou et al., 2011), and A20^-/-^ BMDMs exhibit aberrant spontaneous NLRP3 inflammasome activation, i.e., secretion of mature IL-1β after LPS without ATP stimulation (Walle et al., 2014; Duong et al., 2015). Hence, exaggerated NO produced in A20^tiKO^ BMDMs might drive aberrant NLRP3 inflammasome activation by generating ROS that activate these protein complexes. We measured IL-1β secretion by ELISA, and found that A20^tiKO^ BMDMs secreted IL-1β even in the presence of the iNOS inhibitor 1400w that restored mitochondrial respiration (**Fig. 5G**). Hence, A20 regulates mitochondrial respiration via a distinct cell biological pathway from its regulation of inflammasome activation.

IL-10 deficient macrophages exhibit defects in mitochondrial function after LPS stimulation (Ip et al., 2017). Hence, A20 deficient macrophages might develop mitochondrial dysfunction due to defective IL-10 production. We assayed IL-10 secretion from LPS stimulated A20^tiKO^ and control BMDMs, and found that A20^tiKO^ cells secreted significantly more IL-10 than control cells (**Fig. 5H**). In addition, while mitochondrial defects in IL-10 deficient macrophages do not depend upon NO (Ip et al., 2017), this reactive nitrogen species is critical for compromising mitochondrial respiration in A20 deficient cells. Thus, A20 and IL-10 appear to control mitochondrial function via distinct mechanisms. Nevertheless, the shared metabolic defects of IL-10 deficient and A20 deficient macrophages may underlie overlapping links of these two molecules to early onset inflammatory bowel disease and other inflammatory diseases.

In summary, our findings reveal a profound role for A20 in restraining aerobic glycolysis and supporting metabolic reprogramming toward oxidative phosphorylation. The metabolic impact of A20 on macrophage homeostasis is mediated by iNOS and NO. A20 deficient macrophages share some mitochondrial defects with IL-10 deficient macrophages, but are distinguished by the secretion of high levels of IL-10 and reliance upon iNOS. As A20 deficient macrophages may contribute to inflammatory processes in an expanding array of human diseases, including haploinsufficient A20 patients, understanding the metabolic defects found in these cells should open novel avenues toward controlling these clinical syndromes. Finally, as A20 is a tumor suppressor in both mice (Shao et al., 2013) and humans (Kato et al., 2009; Compagno et al., 2009), the severe skewing toward aerobic glycolysis in A20 deficient macrophages suggests that loss of A20 may also support tumorigenesis via metabolic mechanisms as well as via regulation of cell survival pathways.

## Materials and Methods

### Mice

A20^Flox^ mice were generated in our laboratory and described previously (Tavares et al., 2010). Rosa/ER-Cre mice and iNOS^-/-^ mice were purchased from Jackson Laboratories. All animal research was approved by University of California San Francisco (UCSF) Office of Ethics and Compliance.

### Generation of bone marrow derived macrophages and treatments

Bone marrow derived macrophages were prepared as previously described (Duong et al., 2015). Deletion of A20 *in vitro* was performed by adding 60nM 4-hydroxytamoxifen (4-OHT) 2 days before harvested. The doses of ligands used were as follows: lipopolysaccharide 100 ng/ml; poly(I:C) HMW 20 μg/ml; Pam_3_CSK_4_ 1 μg/ml; CpG-A 2 μM; CpG-B 250 nM, Gardiquimod 1 μg/ml; PolyU (complexed with lipofectamine) 0.5 μg/ml and flagellin (from S. *typhimurium*) 2 μg/ml (Invivogen). For pharmacological inhibitors: allopurinol 25 μM and 1400w 100 μM (Sigma). For macrophage repolarization, cells were stimulated with LPS for 24 hours and washed twice before the addition of 200 ng/ml IL-4 for another 24 hours (PeproTech).

The resulting cells were analyzed by flow cytometry (BD Fortessa) at the UCSF Parnassus Flow Core, RRID:SCR_01826, and the software FlowJo (BD).

### NO, ROS and lactate assays

Culture supernatants from stimulated macrophages were briefly centrifugated to remove any cell debris. Nitric oxide (NO) was measured using a Total Nitric Oxide Assay kit (Invitrogen). To measure intracellular ROS/RNS, macrophages were loaded with 25 μM H_2_DCFDA (Sigma) in serum-free DMEM medium and incubated for 30 min at 37 °C. Cells were then collected, washed once with PBS and resuspended in FACS buffer (2.5% FBS, 0.1% sodium azide, 0.02% sodium bicarbonate in PBS) supplemented with 7-AAD at a working concentration of 2.5 μg/ml for the exclusion of dead cells, and analyzed by flow cytometry (BD Fortessa) at the UCSF Parnassus Flow Core, RRID:SCR_01826, and the software FlowJo (BD). Lactate secretion was measured using a colorimetric Lactate Assay kit (BioVision). All assays were performed according to manufacturer’s instructions.

### Mitochondrial assays

Mitochondrial membrane potential was measured using tetramethylrhodamine methyl ester (TMRM) (Invitrogen) according to manufacturer’s instruction. TMRM (20 nM) was added and incubated for 30 minutes in 37°C CO_2_ incubator before cell harvest. Mitochondrial turnover was examined by first labeling mitochondria in unstimulated cells with MitoTracker Green FM (MTG) (Invitrogen) for 30 minutes according to manufacturer’s instructions. Cells were then washed twice with PBS and stimulated with LPS for 24 hours. MitoTracker Red CM-H_2_Xros

(MTR) (Invitrogen) was then added to the culture 30 minutes before harvest in 37°C CO_2_ incubator. Dead cells were excluded with LIVE/DEAD Fixable Near-IR Viability Dye (Invitrogen) according to manufacturer’s instructions. Fluorescent signals were analyzed with flow cytometry (BD Fortessa) at the UCSF Parnassus Flow Core, RRID:SCR_01826, and the software FlowJo (BD). Mitochondrial DNA (mDNA) to nuclear DNA (nDNA) ration was determined as described in details by Malik and colleagues (Malik et al., 2016) using Quantifast SYBR Master Mix (Qiagen) and analyzed with quantitative real-time PCR (Quantstudio 6 Flex Real-Time PCR Systems from Thermo). To measure the mitochondrial protein Tomm20, cell palettes were subjected to lysis in 1x RIPA lysis buffer (Alfa Aesar) supplemented with complete EDTA-Free Protease Inhibitor Cocktail (Roche), phosphatase inhibitors (1 mM Na_3_VO_4_ and 10 mM NaF) and 10 mM N-ethylmaleimide (Sigma), and centrifuged for 20 minutes (21,000 x g at 4°C). Mitochondrial citrate synthase activity was quantitated using the Citrate Synthase Activity Assay kit (BioVision) according to the manufacturer’s instruction. To measure mitochondrial derived superoxide, the superoxide indicator MitoSOX Red (Invitrogen) was used according to manufacturer’s instructions. Mitochondrial TCA cycle metabolites, GSH and glycine were measured via mass spectrometry as previously described (Louie et al, 2016; Kohnz et al, 2016).

### Quantitative real-time PCR

Total RNA was extracted from BMDMs after treatment using the RNeasy Mini kit (Qiagen) according to the manufacturer’s instructions with on-column DNase I digestion (Qiagen). cDNA was synthesized using the PrimeScript RT-PCR kit (Takara Bio) according to manufacturer’s instructions. Quantitative real-time PCR was performed using TaqMan probes on Quantstudio 6 Flex Real-Time PCR Systems from Thermo.

### Seahorse assays

BMDMs were seeded 2 hours before stimulation on the 24-well Seahorse cell culture plates (Agilent). Cells were stimulated for 24 hours in 37°C CO_2_ incubator, after which glycolytic activity was analyzed using the XF Glycolytic Rate Assay Kit (Agilent). For mitochondrial oxygen consumption, the XF Cell Mito Stress Test Kit was used (Agilent). Both tests were performed according to manufacturer’s instructions.

### Oil Red O staining

BMDMs were seeded on Lab-Tek II 8-well Chamber Slides (Thermo) and stimulated 2 hours after attachment. Cells were harvested by washing in PBS twice and then fixed in 10% NBF (Sigma) at room temperature for 10 minutes. Fixative was removed and fresh 10% NBF was added and allow fixation for another 3 hours. After fixation, cells were washed with ddH_2_O twice and subsequently with 60% isopropanol for 5 minutes at room temperature. The samples were left to dry completely overnight. The next day, samples were stained and incubated with Oil Red O solution (0.21% Oil Red O powder, Sigma, dissolved in isopropanol and filtered, let set at room temperature for 20 minutes before use) for 10 minutes at room temperature. Afterwards the dye was removed and ddH_2_O immediately added. Cells were washed 4 times with ddH_2_O. The slides were allowed to dry and mounted in ProLong Gold Antifade Mountant (Invitrogen) before acquiring images under the microscope (Zeiss Axioscope).

## Acknowledgments

This work was supported by NIH RO1 grants, the UCSF Nutrition and Obesity Research Center (P30DK098722) and the UCSF Liver Center (P30DK026743) from the National Institute of Diabetes and Digestive and Kidney Diseases. Author contributions: M. Chan, B. Malynn, A. Ma conceptualized the study, designed and interpreted the experiments, and wrote the manuscript. M. Chan performed and analyzed the experiments. E. Grossman and D. Nomura performed and analyzed metabolomic experiments. R. Advincula and P. Achacoso assisted with mouse breeding and experiments. The authors declare no competing financial interests.

