## Supplemental Figure 1 for "A20 restriction of nitric oxide production restores macrophage bioenergetic balance"

Supplementary Figure 1: Restoration of M2 repolarization by iNOS inhibitör.
Flow cytometry analysis of CD206 (M2 marker) expression on A20$^{tiKO}$ and WT BMDMs after 24 hours of LPS or LPS+1400w, followed by 24 hours of IL-4.
Note reduced induction of CD206 expression by IL-4 in A20$^{tiKO}$ BMDMs compared with WT cells, and normalization of CD206 expression in A20$^{tiKO}$ BMDMs with iNOS inhibitor 1400w.
Data are representative of two independent experiments.

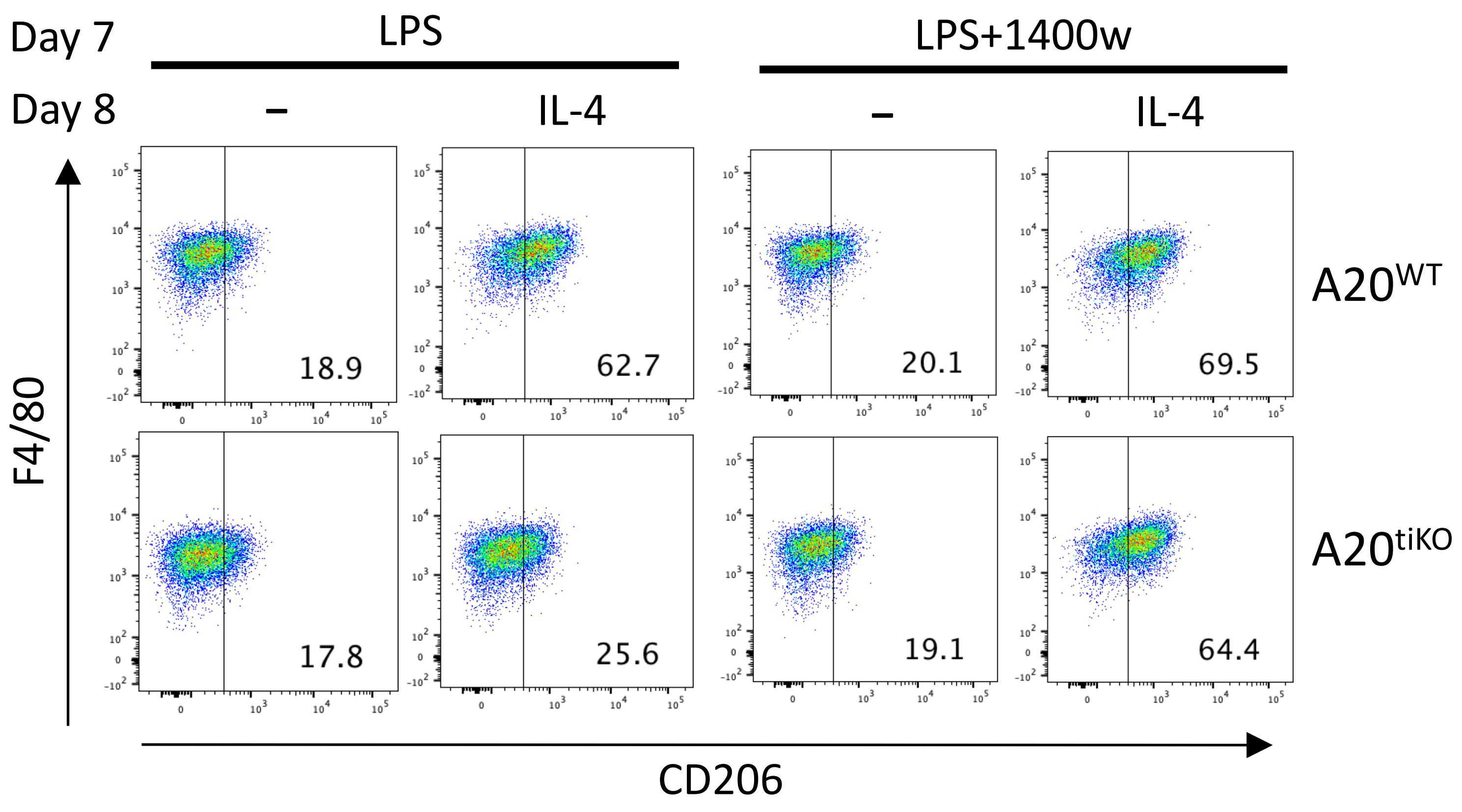
