## Supplemental Figure 2 for "A20 restriction of nitric oxide production restores macrophage bioenergetic balance"

Supplementary Figure 2: Immunoblot of iNOS and A20 protein expression in BMDMs from indicated genotypes. Tubulin expression show below as loading control. Data are representative of three independent experiments.

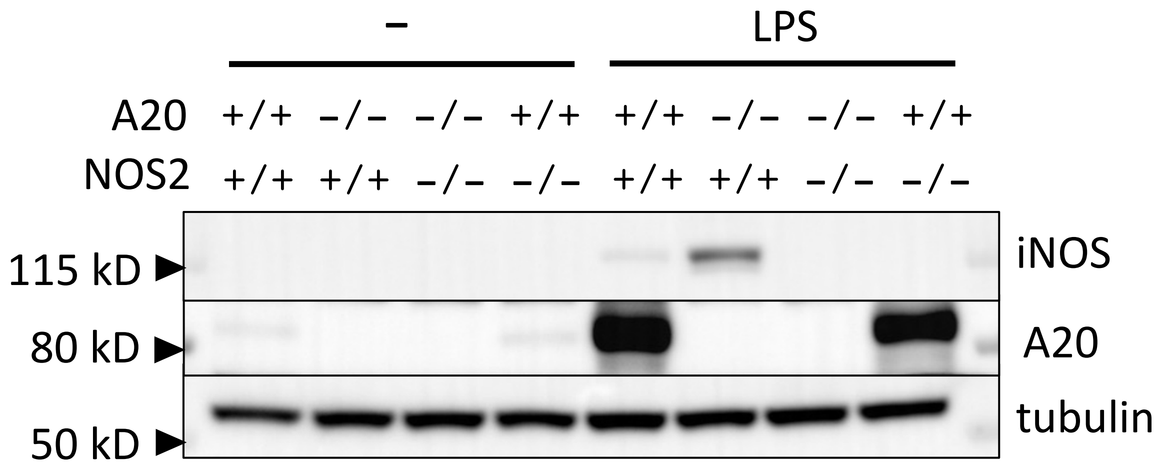
