## Supplemental Figure 3 for "A20 restriction of nitric oxide production restores macrophage bioenergetic balance"

Supplementary Figure 3: Flow cytometric measurement of ROS/RNS in indicated genotypes after 1400w iNOS inhibitor treatment at indicated times.

Note that iNOS inhibition normalizes ROS/RNS in LLPS stimulated A20tiKO cells.

Mean values ± SD are shown. **p<0.01 by unpaired two-tailed t test was used.

Data are representative of three independent experiments.

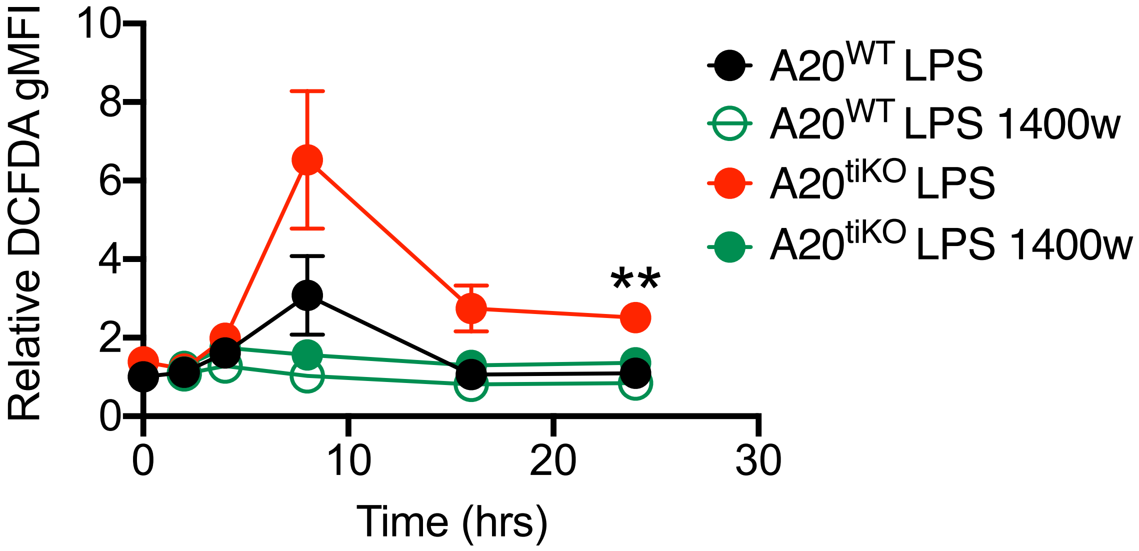
